# Persistent cognitive deficits in anti-LGI1 encephalitis are linked to a reorganization of structural brain networks

**DOI:** 10.1101/2024.03.07.583948

**Authors:** Stephan Krohn, Leonie Müller-Jensen, Joseph Kuchling, Amy Romanello, Thorsten Bartsch, Frank Leypoldt, Friedemann Paul, Harald Prüss, Carsten Finke

## Abstract

**Importance:** Despite immunotherapy, most patients with anti–leucine-rich, glioma-inactivated 1 encephalitis (LGI1-E) develop long-term cognitive deficits that persist for years after peak illness. However, the structural brain changes that underlie these deficits remain poorly understood.

**Objective:** To study the relationship between cognitive outcomes and white matter (WM) networks in LGI1-E.

**Design:** Cross-sectional study.

**Setting:** German university center (Charité - Universitätsmedizin Berlin).

**Participants:** 25 patients with LGI1-E (19/25 male [76%], mean age: 63 ± 12 years) and 25 age- and sex-matched healthy controls (HC), recruited between January 2013 and April 2019.

**Main Outcomes and Measures:** Clinical assessments including the modified Rankin Scale (mRS) and Clinical Assessment Scale in Autoimmune Encephalitis (CASE); comprehensive cognitive testing; WM tractography using diffusion-weighted MRI.

**Results:** All patients had received first-line immunotherapy, and two-thirds underwent second-line immunotherapy. Patients showed a significant reduction in mRS scores from peak illness to post-acute follow-up (*z* = -3.8, *p* < 0.001, n = 20), with 85% presenting “good” functional outcomes (post-acute mRS ≤ 2), paralleled by a significant reduction in CASE scores (*z* = -3.5, *p* < 0.001, n = 20).

Despite this overall improvement, however, cognitive symptoms were highly prevalent at peak illness (95% of patients affected) and strongly persisted into the post-acute disease stage (85% affected).

Neuroimaging at post-acute follow-up (median: 12 months from onset) revealed that LGI1-E is characterized by (i) significantly reduced whole-brain structural connectivity (*t* = -2.16, *p* = 0.036, *d* = -0.61), (ii) a cortico-subcortical hypoconnectivity cluster that strongly affects the hippocampus but also severely impacts extra-limbic brain systems, (iii) systematic limbic and extra-limbic decreases in node degree — a graph-theoretical measure of overall connectedness, and (iv) a “topological reorganization” of structural brain networks, marked by a bidirectional shift in the relative importance of individual brain regions in the network.

Importantly, the extent of this network reorganization was significantly related to persistent cognitive deficits in the domains of verbal memory (*r* = -0.57, *p* = 0.007, n = 21), attention (*r* = -0.47, *p* = 0.030, n = 21), and executive functions (*r* = -0.60, *p* = 0.010, n = 17).

**Conclusion and Relevance:** This study characterizes LGI1-E as a network disease that affects both limbic and extra-limbic brain systems and shows that a reorganization of WM networks is linked to multi-domain cognitive deficits in the post-acute disease stage – despite immunotherapy and good overall recovery. These findings highlight the need for extended treatment strategies to improve long-term cognitive outcomes and propose a sensitive new neuroimaging marker to include in prospective clinical trials.

**Key Points:** *Question:* What structural brain changes underlie the persistent cognitive deficits observed in patients with anti–leucine-rich, glioma-inactivated 1 encephalitis (LGI1-E)?

*Findings:* This cross-sectional study shows that LGI1-E is characterized by a structural reorganization of white matter networks that affects both limbic and extra-limbic brain systems and correlates with persistent deficits in verbal memory, attention, and executive functions at post-acute follow-up – despite immunotherapy and good overall clinical recovery.

*Meaning:* This study characterizes LGI1-E as a network disease –beyond focal damage to the limbic system– and shows that persistent cognitive deficits relate to immunotherapy-resistant changes in structural brain networks, highlighting the need for extended treatment strategies to improve long-term cognitive outcomes.

## Introduction

Anti–leucine-rich, glioma-inactivated 1 encephalitis (LGI1-E) represents the most common form of autoimmune encephalitis (AE) in older adults.^1^ Caused by autoantibodies against the LGI1 neuronal surface antigen,^2^ LGI1-E commonly presents with prodromal symptoms including the pathognomonic faciobrachial dystonic seizures (FBDS) that can precede other manifestations by weeks to months.^3–5^ Subsequently, most patients develop a combination of memory impairment, temporal lobe seizures, sleep disturbances, confusion, and behavioral abnormalities^1, 6^ – a clinical syndrome conceptualized as “limbic encephalitis”^7, 8^ that is frequently accompanied by T2/FLAIR hyperintensities of the medial temporal lobe (MTL) in clinical magnetic resonance imaging (MRI).^9, 10^

More recently, however, advanced neuroimaging studies have suggested a key role of extra-limbic brain systems in LGI1-E, complemented by clinical evidence on long-term outcomes showing that our understanding of LGI1-E remains far from complete.

Specifically, imaging studies have demonstrated that LGI1-E is characterized by a widespread disruption of functional brain networks beyond the limbic system, including the default mode, sensorimotor, salience, and higher visual network,^11, 12^ paralleled by metabolic abnormalities in the basal ganglia, motor areas, and prefrontal cortex.^13–16^ Similarly, microstructural brain damage as assessed with diffusion-weighted imaging (DWI) has been observed in both limbic and extra-limbic brain structures and correlates with poorer functional outcome.^9, 17^

Importantly, clinical studies have identified cognitive deficits as a crucial long-term outcome of LGI1-E, showing that these deficits (i) persist for years after peak illness,^5, 9, 18, 19^ (ii) encompass diverse cognitive functions beyond isolated memory impairment,^18–20^ and (iii) represent a key predictor of long-term functional disability.^21^ However, the structural brain changes that underlie these deficits remain poorly understood.

Here we hypothesized that –beyond focal damage to individual regions– LGI1-E specifically affects the brain’s white matter (WM) network that connects limbic and extra-limbic brain structures. Therefore, we combined DWI-based probabilistic tractography,^22^ graph-theoretical analyses,^23, 24^ clinical evaluation, and comprehensive neuropsychological assessments to study the relationship between WM networks and cognitive outcomes in patients with LGI1-E.

## Methods

### Study cohort

This study included 25 patients with LGI1-E recruited from a German university referral center (Charité - Universitätsmedizin Berlin) and 25 healthy control participants (HC) matched for age and sex. All patients fulfilled current diagnostic criteria^25^ and tested positive for LGI1 antibodies in serum and/or CSF. The study was approved by the local ethics committee, and all participants gave written informed consent. For further details on data collection and all methodological approaches below, please refer to the Extended Methods in the Supplement.

### MRI Analyses

WM networks were estimated with an optimized tractography protocol,^26^ as recently validated in AE.^27^ Briefly, DWI processing was performed with MRtrix3,^28^ FSL,^29^ and ANTs,^30^ including denoising, eddy-current correction, motion correction, and global intensity normalization using an average response function across the study cohort. Anatomical T1-weighted scans were parcellated into 84 gray matter (GM) regions using the Desikan-Killiany atlas^31^ to enable anatomically constrained probabilistic tractography and spherical-deconvolution informed filtering of tractograms with 2×10^7^ streamlines and inter-individual connection density normalization.^26, 32–34^

The resulting WM networks were analyzed regarding structural connectivity (SC) and subjected to graph-theoretical analyses of network topology (i.e., the organizational structure of how individual GM regions are connected to each other), using the Brain Connectivity Toolbox.^35^ Therein, ‘nodes’ in the network correspond to GM regions and ‘edges’ to WM tracts, allowing for an estimation of node degree as a measure of a region’s overall connectedness and betweenness centrality as a measure of relative importance in the network.^23^

Anatomical regions were assigned to functional brain systems using a new region-to-network mapping procedure, and alterations of network organization were quantified with a novel “Topology Deviation Index” (TDI), as detailed in the Supplement.

### Clinical and cognitive outcomes

Following previous approaches,^11^ we applied a standardized neuropsychological test battery to quantify cognitive performance with composite *z*-scores across the domains of visuospatial memory, verbal episodic memory, attention, executive functions, and working memory, where higher scores indicate better performance. Functional neurological outcomes were rated retrospectively with the modified Rankin Scale (mRS) and the Clinical Assessment Scale in Autoimmune Encephalitis (CASE).^36^

### Statistical Analysis

Univariate group differences were assessed with independent-sample t-tests, ordinal paired outcomes scores with signed rank tests, continuous relationships with product-moment or rank-based correlation, and imaging markers with permutation tests. When a variable was repeatedly tested against multiple other measures, we applied the Benjamini-Hochberg procedure to control the false discovery rate (FDR).^37^

## Results

### Clinical features

The study population included 25 patients with LGI1-E (6 females; mean age: 63 ± 12 years) and 25 HC (10 females; 59 ± 11 years) matched for age (t = 1.38, p = 0.175) and sex (χ^2^ = 0.83, p = 0.363). All patients had received first-line immunotherapy (IV steroids, plasma exchange, IV immunoglobulins), and 68% underwent second-line immunotherapy (rituximab, azathioprine). Clinical characteristics of the patient sample are summarized in Table 1.

**Table 1.** Clinical features of the patient sample.

| Characteristic | Patients, no. (%) |
| --- | --- |
|  | LGI1-E (n = 25) |
| **Symptoms in acute disease stage** |  |
| Memory impairment | 24 / 25 (96) |
| Psychiatric symptoms | 14 / 25 (56) |
| Sleep disturbance | 8 / 22 (36) |
| FBDS | 12 / 25 (48) |
| Other seizures (including autonomic seizures) | 16 / 25 (64) |
| SIADH / hyponatremia | 13 / 25 (52) |
| **Diagnostics** |  |
| **MRI** |  |
| Any MRI abnormality* | 17 / 23 (74) |
| Hippocampal T2/FLAIR hyperintensity / edema (bilaterally) | 6 / 23 (26) |
| Hippocampal T2/FLAIR hyperintensity / edema (unilaterally) | 5 / 23 (22) |
| Basal ganglia T2/FLAIR hyperintensity / edema | 1 / 23 (4) |
| **EEG abnormalities\*\*** | 12 / 21 (57) |
| **Malignant tumor disease** | 0 / 22 (0) |
| **Treatment** |  |
| Days from onset to treatment, median (IQR) | 48 (11-133) |
| Steroids (IV) | 25 / 25 (100) |
| Plasma exchange | 16 / 25 (64) |
| Intravenous immunoglobulins (IVIG) | 9 / 25 (36) |
| Rituximab | 14 / 25 (56) |
| Azathioprine | 5 / 25 (20) |
| Antipsychotic or antidepressant medication | 3 / 25 (12) |
| Antiseizure medication | 20 / 25 (80) |
| **Months from onset to MRI, median (IQR)** | 12 (6-23) |
| **Outcome** |  |
| mRS score at peak (median, range) | 3 (2-4) |
| mRS score at MRI (median, range) | 1 (0-3) |
| CASE score at peak (median, range) | 4 (2-8) |
| CASE score at MRI (median, range) | 2 (0-5) |
| Relapse | 7 / 19 (37) |
| Fatal outcome | 0 / 25 (0) |

\* Includes: Medial temporal lobe T2/FLAIR hyperintensity and edema, unilateral or bilateral hippocampal T2/FLAIR hyperintensity with or without gadolinium enhancement, global atrophy, medial temporal lobe atrophy, white matter lesions, leukoencephalopathy.
\*\* Includes: Diffuse slow activity, focal slow activity, epileptic activity.
CASE, Clinical Assessment Scale for Autoimmune Encephalitis; CSF, cerebrospinal fluid; FBDS, faciobrachial dystonic seizures; FLAIR, fluid-attenuated inversion recovery; mRS, modified Rankin Scale; SD, standard deviation; SIADH, syndrome of inappropriate antidiuretic hormone secretion; IVIG intravenous immunoglobulins.

### Structural connectivity analysis

First, we assessed whether patients with LGI1-E exhibit systematic differences in structural connectivity (SC) estimates. To this end, we extracted the unthresholded structural connectomes and the total number of estimated tracts for each participant to compare region-to-region SC (**Fig. 1A**) and whole-brain SC (**Fig. 1B**) between patients and HC. Here, patients predominantly presented reduced SC among individual GM regions (**Fig. 1A**) and showed significantly lower whole-brain SC compared to HC (**Fig. 1B**; *t* = −2.16, *p* = 0.036, *d* = −0.61).

**Fig. 1.**
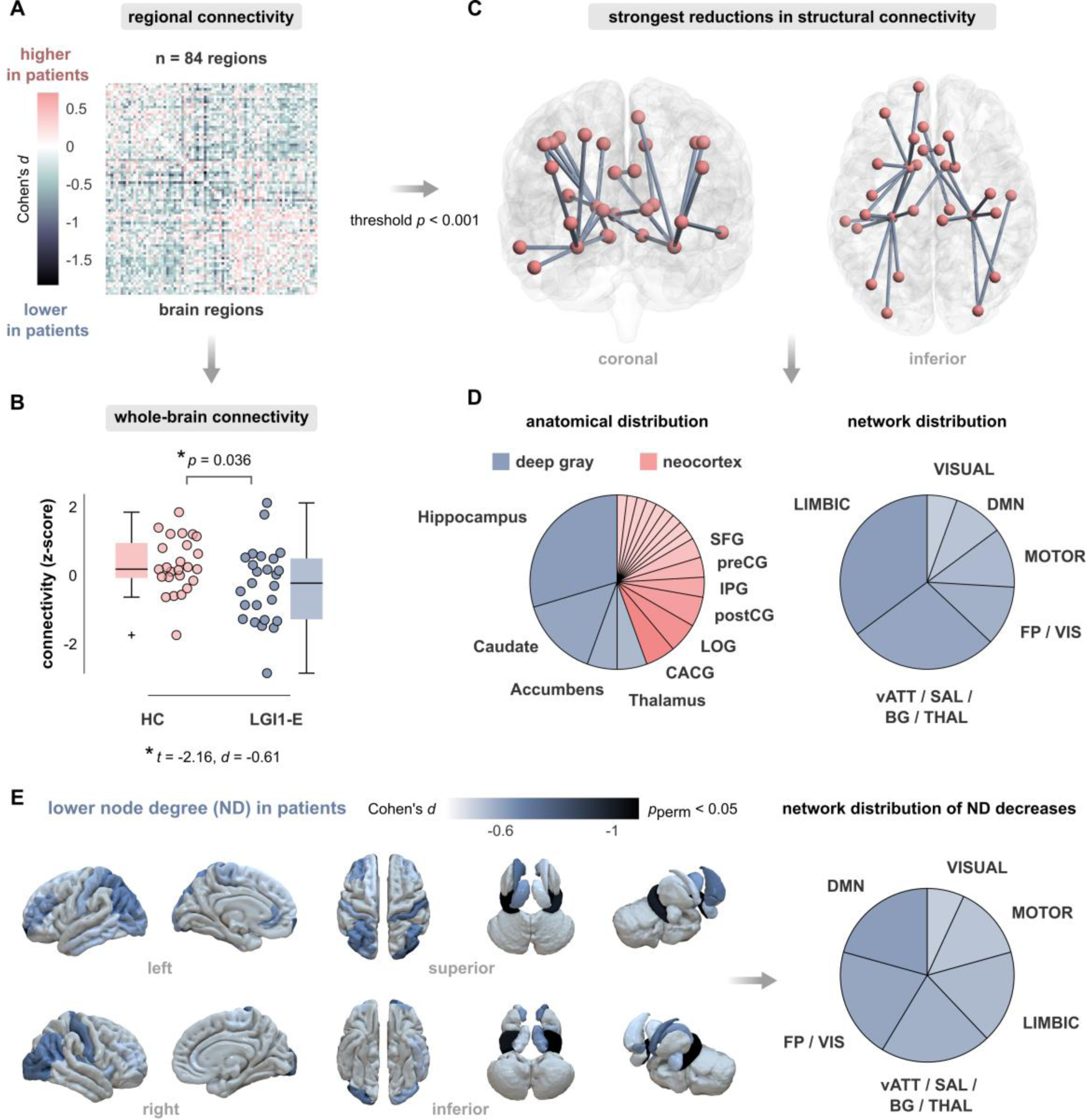
Structural brain networks in anti-LGI1 encephalitis are characterized by a cortico-subcortical hypoconnectivity cluster beyond the limbic system. **(A)**, Differences of structural connectivity strength between patients with anti-LGI1 encephalitis (LGI1-E) and healthy control participants (HC) across all 84 brain regions. The color bar represents the connectome-wide effect sizes of between-group comparisons (Cohen’s *d*, n = 25 participants each). **(B)**, Patients with LGI1-E show a reduction in whole-brain structural connectivity strength compared to HC. **(C)**, Network representation of region-wise connectivity differences from panel (A) in brain space. Group differences are thresholded to *p*<0.001, at which only negative effects remain (i.e., reduced connectivity in LGI1-E). Nodes represent brain regions, and edges represent reduced structural connectivity between them. **(D)**, Distribution of the connectivity reductions from panel (C) over anatomical structures (left) and functional network assignments (right). Anatomical labels correspond to the Desikan-Kiliany atlas.^31^ For neocortical areas, gyri are abbreviated as follows: CACG: caudal anterior cingulate, IPG: inferior parietal, LOG: lateral occipital, postCG: postcentral, preCG: precentral, and SFG: superior frontal gyrus. Cortical regions with only one entry are omitted for visibility and correspond to the following areas: superior temporal, middle temporal, inferior temporal, rostral anterior cingulate, paracentral, superior parietal, rostral middle frontal, caudal middle frontal, and frontal pole. Functional assignments rest on a novel region-to-network mapping procedure (see Supplement). Abbreviations – DMN: default mode network; FP/VIS: fronto-parietal network/visual downstream; LIMBIC: mesolimbic network; MOTOR: somatomotor network; vATT/SAL/BG/THAL: ventral attention network/salience network/basal ganglia/thalamus; VISUAL: visual network. **(E)**, Brain areas exhibiting decreased node degree (ND) in patients with LGI1-E. The color scale maps the effect size of the comparison to HC given by Cohen’s *d* (LGI1-E vs. HC, n = 25 each; threshold: permuted p < 0.05). The right subpanel visualizes the distribution of brain areas with lower ND over functional systems.

To assess the spatial distribution of these SC reductions, we subsequently thresholded the network-wide differences to *p* < 0.001 and observed the strongest effects in a spatial cluster of deep GM structures and neocortical areas (**Fig. 1C**). Specifically, these connectivity reductions clustered in the hippocampus, caudate, accumbens, and thalamus as well as a variety of neocortical areas, most notably the caudal anterior cingulate, lateral occipital gyrus, the pre- and postcentral gyri and the inferior parietal and superior frontal gyri (**Fig. 1D**, *left*). To estimate how this cortico-subcortical hypoconnectivity cluster relates to functional brain systems, we applied a novel region-to-network mapping procedure using the 7-system Multiresolution Intrinsic Segmentation Template (see Supplement).^38^ This approach revealed that connectivity reductions clustered in the mesolimbic network – in line with the traditional conception of LGI1-E as a form of ‘limbic’ encephalitis^7, 8^ – but additionally affected a wide range of extra-limbic brain systems including attentional / salience, motor, visual, and default mode areas (**Fig. 1D**, *right*).

Further supporting these findings, graph analysis of structural connectomes showed that LGI1-E is characterized by systematic decreases in node degree (ND) – a measure that quantifies the overall connectedness of individual brain regions (see Supplement). Here again, we found ND to be most strongly decreased in the hippocampus (left: *d* = −1.16, *p*_perm_ < 0.001; right: *d* = −1.01, *p*_perm_ < 0.001), but significant reductions were also observed across multiple other subcortical and neocortical areas (**Fig. 1E**, *left*). As for raw connectivity estimates, network mapping revealed widespread extra-limbic ND reductions, most notably including default mode, fronto-parietal, and attentional / salience areas (**Fig. 1E**, *right*).

### Network topology analysis

Given these widespread SC alterations, we next asked if LGI1-E also affects the topological organization of structural brain networks beyond connectivity reductions. To this end, we applied a graph-theoretical framework and estimated the normalized betweenness centrality (BC) for each brain region. Briefly, BC measures the relative importance of a node in the network by quantifying the number of all shortest paths in the network that pass through it. In consequence, a brain region shows high BC if it lies on many shortest paths connecting other brain regions and thus exerts a ‘bridging role’ in the network.^39^ Comparing BC values between LGI1-E and HC revealed that patients exhibit bidirectional changes in network centrality:

**Figure 2A** shows that BC was *increased* in patients in the amygdala bilaterally (left: *d* = 0.61, *p*_perm_ = 0.016; right: *d* = 0.72, *p*_perm_ = 0.006) and across various neocortical regions (strongest effects: left inferior parietal cortex: *d* = 0.63, *p*_perm_ = 0.010; left parahippocampal gyrus: *d* = 0.61, *p*_perm_ = 0.014). These BC increases clustered in multiple brain areas beyond the limbic system, most prominently the fronto-parietal network (**Fig. 2A**, *right*).

**Fig. 2.**
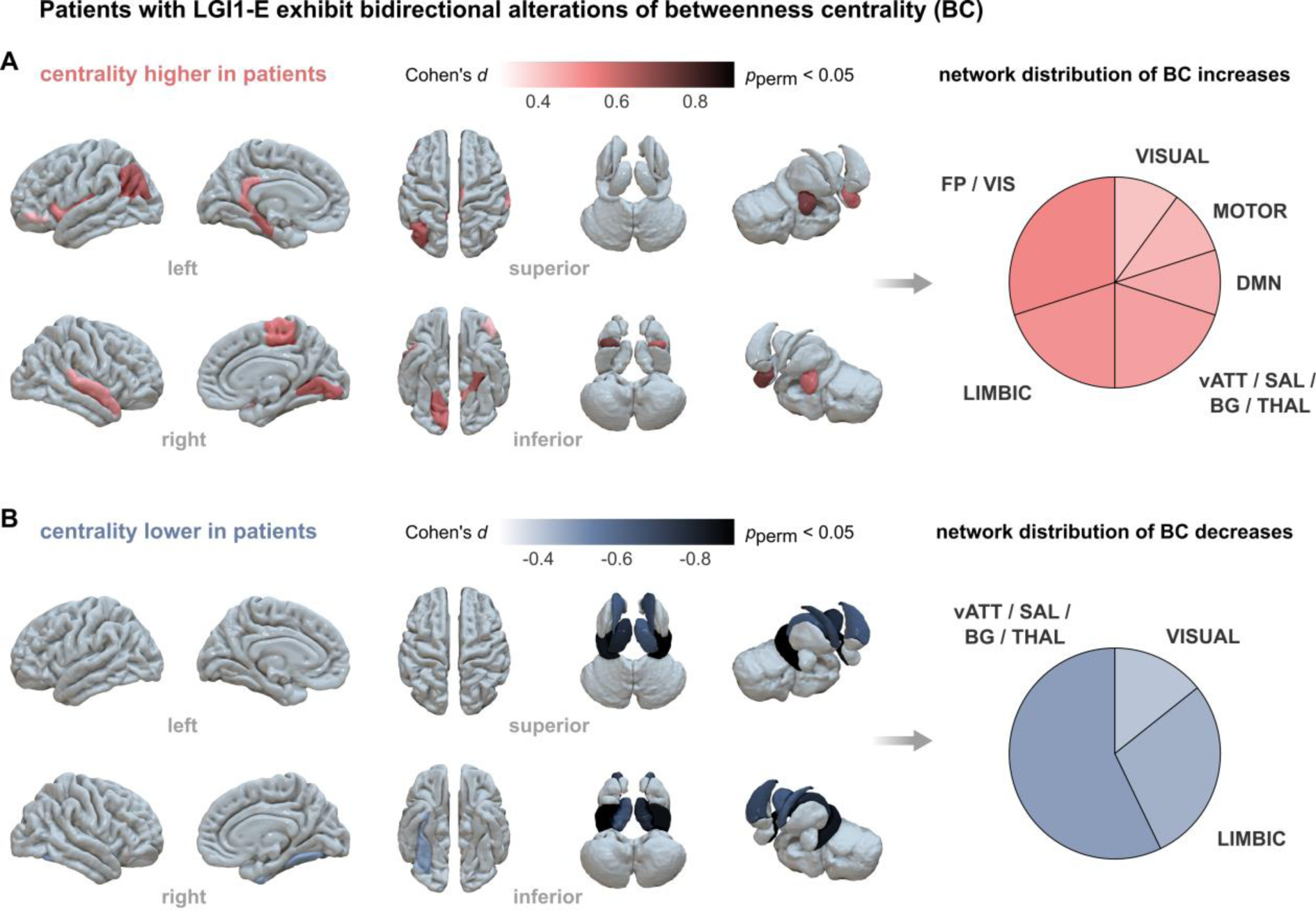
Patients with anti-LGI1 encephalitis exhibit bidirectional changes of structural network topology. **(A)**, Brain areas exhibiting increased betweenness centrality (BC) in patients with anti-LGI1 encephalitis (LGI1-E). The color scale maps the effect size of the comparison to healthy control (HC) participants, given by Cohen’s *d* (LGI1-E vs. HC, n = 25 each; threshold: permuted *p* < 0.05). BC increases primarily affect cortical areas and the amygdala bilaterally. The right panel visualizes the distribution of BC increases in region-to-network mapping. **(B)**, Brain areas exhibiting decreased BC in patients with LGI1-E. BC decreases primarily affect the basal ganglia, the thalamus, and the hippocampus bilaterally. The color scale maps the effect size of the between-group comparison, and the right panel visualizes the network distribution of BC decreases. Functional assignments rest on region-to-network mapping (see Supplement). Abbreviations – DMN: default mode network; FP/VIS: fronto-parietal network/visual downstream; LIMBIC: mesolimbic network; MOTOR: somatomotor network; vATT/SAL/BG/THAL: ventral attention network/salience network/basal ganglia/thalamus; VISUAL: visual network.

Simultaneously, however, patients with LGI1-E also showed *decreased* BC across several other brain regions (**Fig. 2B**). These centrality reductions clustered in deep GM areas and were strongest in the hippocampal formations (right: *d* = −0.96, *p*_perm_ < 0.001; left: *d* = −0.84, *p*_perm_ = 0.002), the right caudate nucleus (*d* = −0.71, *p*_perm_ = 0.006), and the left thalamus (*d* = −0.78, *p*_perm_ = 0.005). Here again, extra-limbic brain systems were strongly affected, most notably subcortical and attentional / salience areas (**Fig. 2B**, *right*).

In sum, patients with LGI1-E showed a “topological reorganization” of structural brain networks, which is characterized by a bidirectional shift in the relative importance of individual brain regions in the network.

### Clinical outcomes

Next, we sought to characterize the clinical outcomes in our study cohort from peak illness to the post-acute study visits at which the MRI recordings were acquired (median time from onset to MRI: 12 months, IQR 6-23).

Therefore, we analyzed clinical scores on the modified Rankin Scale (mRS) and the more disease-specific Clinical Assessment Scale in Autoimmune Encephalitis (CASE). Patients showed a significant reduction in mRS scores from peak to post-acute follow-up (**Fig. 3A**; *z* = −3.8, *p* < 0.001, n = 20), with 85% of patients presenting a post-acute mRS ≤ 2, commonly regarded as a “good” functional outcome.^40^ Similarly, the CASE sum score across all clinical subdomains decreased markedly from peak to post-acute follow-up (**Fig. 3B**; *z* = −3.5, *p* < 0.001, n = 20). However, patient outcomes varied strongly across the CASE subdomains (**Fig. 3C**, *left*), with memory dysfunction being the most prevalently affected domain at peak illness (95% of patients affected), followed by seizures (90%), and psychiatric symptoms (70%). Importantly, only 15% of patients showed complete recovery of memory symptoms at post-acute follow-up, while this was 35% for seizures and 55% for psychiatric symptoms (**Fig. 3C**, *right*). In sum, patients thus presented long-term cognitive symptoms despite good overall recovery on standard clinical assessment scales.

**Fig. 3.**
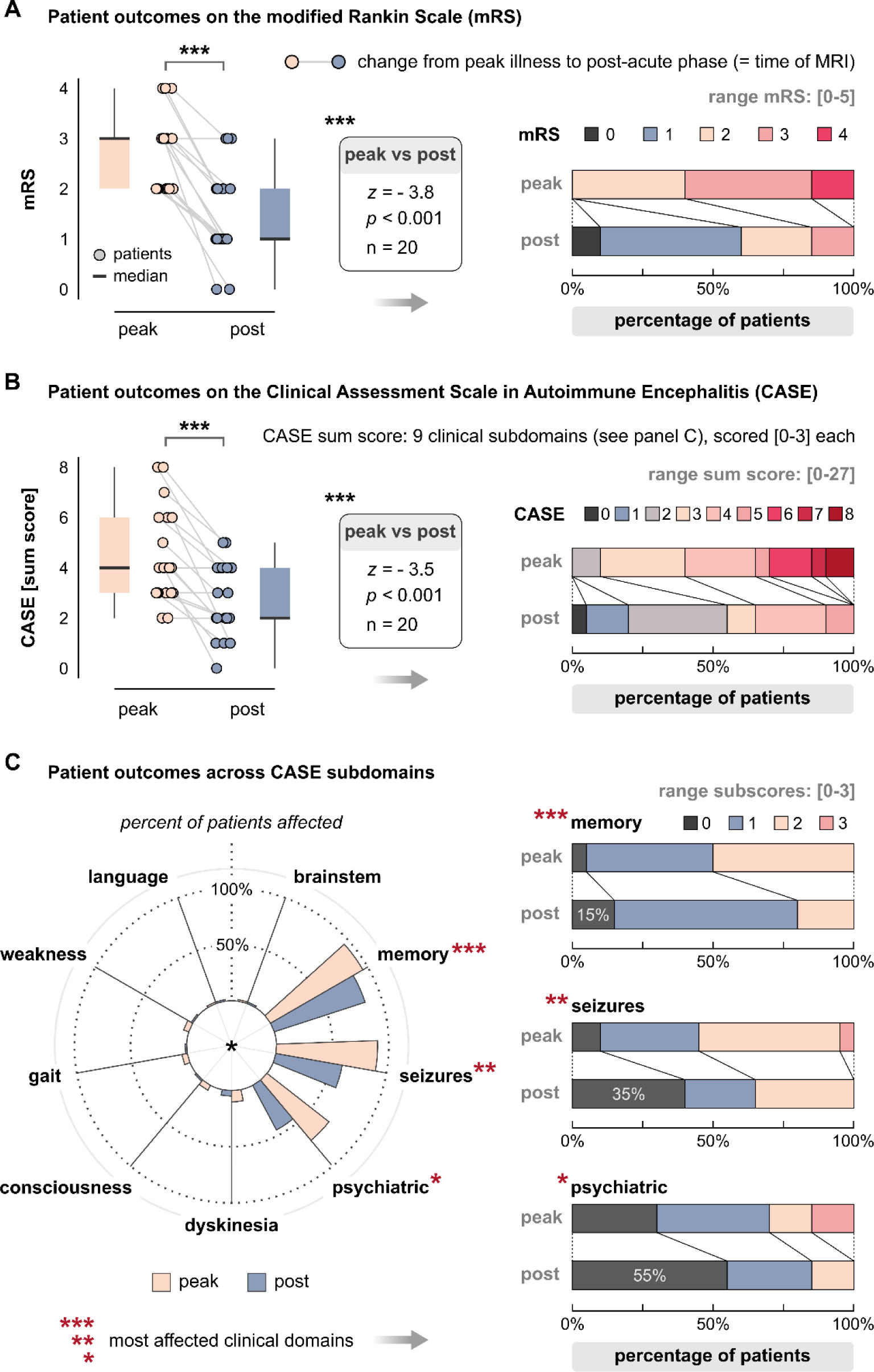
Patients with anti-LGI1 encephalitis show long-term cognitive symptoms despite good overall recovery on standard clinical assessment scales. **(A)**, Patient outcomes on the modified Rankin Scale (mRS) from peak illness to the post-acute phase. MRI recordings underlying the imaging analyses were obtained at post-acute follow-up. Peak vs. post-acute scores were compared with the Wilcoxon signed rank test. **(B)**, Patient outcomes on the Clinical Assessment Scale in Autoimmune Encephalitis (CASE). The CASE sum score is given by the sum over 9 clinical subdomains, each scored with [0-3]. **(C)**, Patient outcomes across CASE subdomains. The polar plot shows the percentage of affected patients at peak illness and post-acute follow-up. The bar plots relate patient outcomes across the three most prevalent clinical subdomains: memory dysfunction, seizures, and psychiatric symptoms.

### Cognitive outcomes

Given this clinical course, we lastly focused on cognitive outcomes specifically and asked how the topological reorganization of structural brain networks relates to post-acute cognitive performance in LGI1-E. To this end, we first computed a reference topology over all HC participants as the median BC for every brain region (**Fig. 4A**). We then defined a topology deviation index (TDI) as the non-parametric correlation distance between this reference distribution in HC and the corresponding centrality values of each individual participant (see Supplement). Figure 4B illustrates this approach for one participant whose individual brain topology closely adheres to the reference (*left panel;* low TDI; a control participant) and another one whose topology deviates strongly from the reference (*right panel*; high TDI; a patient with LGI1-E).

**Fig. 4.**
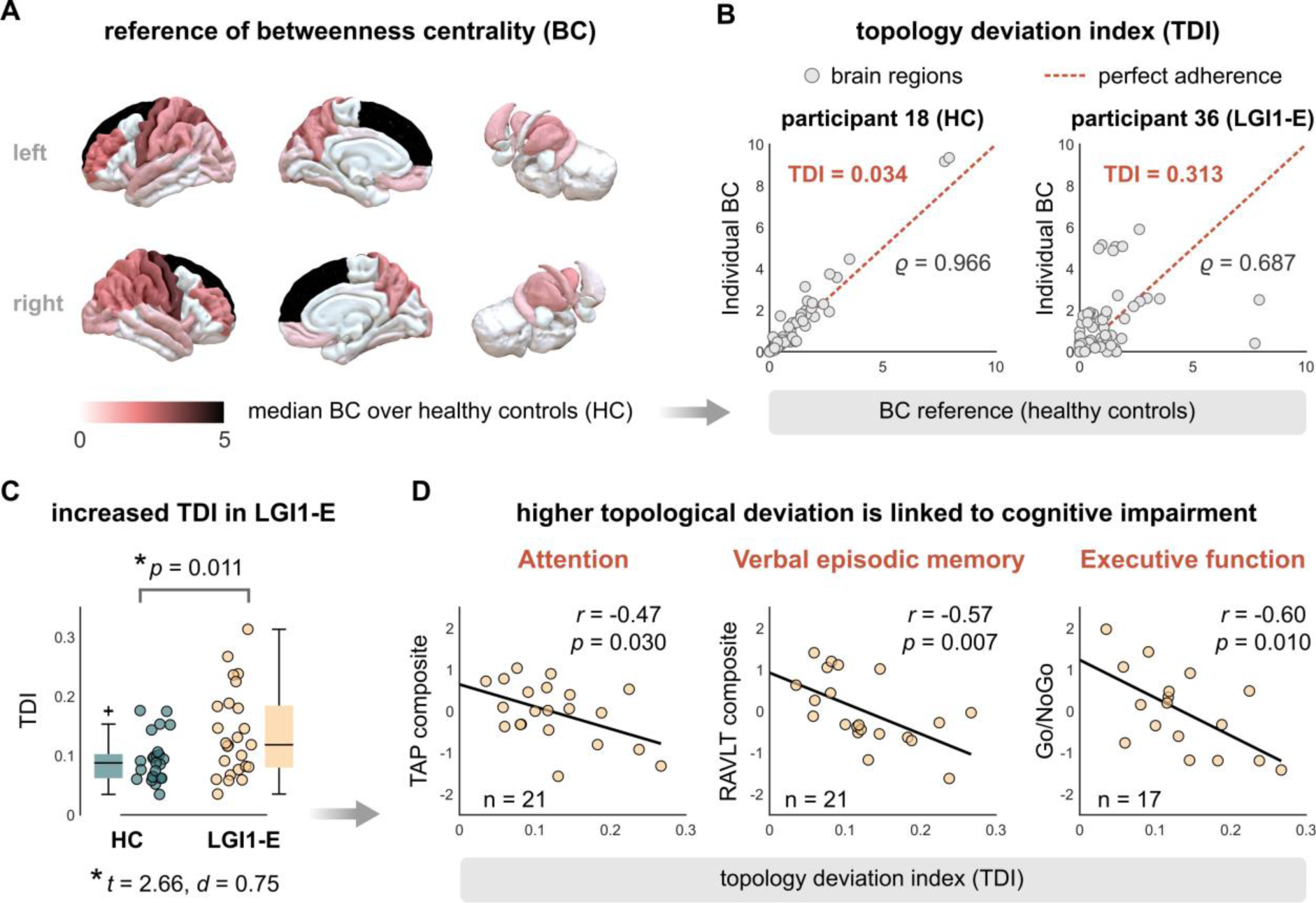
Deviation of structural network topology from healthy reference is associated with multi-domain cognitive impairment in patients with anti-LGI1 encephalitis. **(A)**, Reference distribution of betweenness centrality (BC), computed as the median BC value across all 25 healthy control participants (HC) for every brain region. **(B)**, Illustration of the topology deviation index (TDI), computed as the non-parametric correlation distance between the BC distribution of an individual participant (y-axis) and the reference distribution from panel (A) (x-axis). The left plot illustrates a control participant with low topological deviation, the right plot a patient with high deviation. **(C)**, Patients with anti-LGI1 encephalitis show significantly higher TDI scores than HC (*t*-test, n = 25 each group, *d*: effect size). **(D)**, Higher TDI scores in patients are associated with impaired cognitive performance across multiple cognitive domains.

We then tested the difference in TDI values between HC and patients with LGI1-E. Therein, we hypothesized that the HC group should show lower TDI values because (i) each healthy individual formed part of the group on which the reference was defined and because (ii) patients showed bidirectional alterations of BC in between-group testing. As expected, patients exhibited significantly higher TDI values than HC, with medium to large effect size (*t* = 2.66, *p* = 0.011, *d* = 0.75; **Fig. 4C**). Subsequently, we related the TDI values of individual patients to their cognitive outcomes scores in the domains of working memory, visuospatial memory, verbal episodic memory, executive function, and attention. While we observed no relationship between TDI and working memory (*r* = −0.06, *p* = 0.78, n = 24, *p*_FDR_ = 0.78) or visuospatial memory (*r* = −0.08, *p* = 0.74, n = 20, *p*_FDR_ = 0.78), higher topological deviation in patients was significantly associated with reduced cognitive performance in the domains of verbal episodic memory (*r* = −0.57, *p* = 0.007, n = 21, *p*_FDR_ = 0.026), attention (*r* = −0.47, *p* = 0.030, n = 21, *p*_FDR_ = 0.050), and executive functions (*r* = −0.60, *p* = 0.010, n = 17, *p*_FDR_ = 0.026; **Fig. 4D**).

Finally, exploratory follow-up analyses showed that patients who had received second-line immunotherapy did not differ from those with first-line therapy only – neither in terms of TDI values (*t* = 0.38, *p* = 0.71) nor cognitive composite scores (all *p* > 0.05).

## Discussion

In this cross-sectional study, we combined probabilistic tractography, computational methods, clinical evaluation, and cognitive assessments to study the relationship between structural brain damage and cognitive outcomes in patients with LGI1-E. Specifically, we show that (i) LGI1-E is characterized by structural connectivity reductions that cluster in limbic areas but additionally affect a wide range of extra-limbic brain systems, (ii) structural brain networks in LGI1-E undergo a “topological reorganization” marked by a bidirectional shift in the relative importance of individual brain regions in the network, (iii) patients with LGI1-E show persistent cognitive deficits despite good overall clinical recovery, (iv) these deficits affect multiple cognitive domains beyond memory dysfunction, and (v) this multi-domain cognitive impairment is strongly linked to brain network reorganization in the post-acute disease stage. Collectively, these findings advance our understanding of LGI1-E along three principal directions.

### Cognitive outcomes of anti-LGI1 encephalitis

First, it has become increasingly clear that cognitive deficits represent a key clinical outcome of LGI1-E.^6, 9, 18, 21, 40–44^ On the one hand, recent evidence suggests that cognitive impairment in LGI1-E is not limited to memory deficits^9^, which are frequently observed in limbic encephalitis due to affliction of the medial temporal lobe. Instead, patients with LGI1-E commonly show *multi-domain* deficits beyond memory impairment^18–20^, involving executive functions, language skills, psychomotor speed, and attentional capacities – cognitive domains not classically attributed to the limbic system. Our study clearly supports this view by showing that structural brain changes are not only linked to reduced performance in the domain of verbal episodic memory, but also relate to persistent deficits in executive functions and attention.

On the other hand, there is accumulating evidence that cognitive deficits in LGI1-E persist well into the post-acute disease phase. Although up to 80% of patients show fast responses to immunotherapy with regard to seizures, cognitive recovery is much more protracted, with most patients reporting residual cognitive deficits after ≥ 2 years.^5^ Importantly, even with immunotherapy, only about 35% of patients with LGI1-E return to their baseline cognitive function, and at least 20% of patients require assistance in daily living due to persistent cognitive deficits.^45^ Strikingly, even though all our patients had received immunotherapy at peak illness, we found that cognitive symptoms strongly persisted into the post-acute phase – in spite of significant overall improvement on the mRS and the more disease-specific CASE score.^36^ These findings converge well with recent related reports on long-term outcomes in LGI1-E, which similarly observed significant residual symptoms across cognition and fatigue.^21, 40^ As such, even though patients may show good recovery in terms of mRS or CASE, it has been argued that these improvements are not ‘good enough’^46^ given that at least two thirds of patients suffer from residual deficits of cognition, mood, or fatigue, and only about 15% are able to return to work.^1, 40^

In consequence, a crucial open question is how long-term cognitive deficits in LGI1-E relate to concomitant changes in brain structure. Our study identifies such a link by showing that cognitive performance in the post-acute stage is more impaired in those patients whose brains exhibit a higher degree of structural reorganization. Notably, this relationship is not confined to focal brain regions but rather features many simultaneous changes across the global WM network.

### Anti-LGI1 encephalitis as a network disease

A second open question concerns the spatial pattern of brain damage in LGI1-E. In line with previous studies,^9, 47, 48^ we observed strong alterations in limbic brain structures, and specifically the hippocampus which showed significant reductions in raw structural connectivity, node degree (a graph measure of overall connectedness), and betweenness centrality (a measure of relative importance in the network). Crucially, however, region-to-network mapping revealed that LGI1-E additionally involves widespread alterations of extra-limbic brain systems, including not only the basal ganglia, amygdalae, and thalamus, but also higher-order cognitive systems such as the default-mode network and salience areas. Although LGI1-E is traditionally conceptualized as a form of limbic encephalitis,^7, 8^ these findings suggest it should rather be understood as a “network disease” – a view that may help reconcile diverse previous findings on clinical course and neuroimaging changes.

First, our findings align remarkably well with the widespread disruption of functional brain networks in LGI1-E,^11^ suggesting that these changes in brain activity are a direct expression of alterations in the WM tracts connecting spatially distant GM regions. Similarly, a recent study has suggested that memory impairment in AE may not be entirely explained by hippocampal dysfunction alone but rather by wider network alterations involving the cingulate, thalamus, precuneus, prefrontal cortex, and posteromedial cortex.^11, 49^ As abnormalities in clinical MRI are typically strongest in the MTL —where LGI1 expression clusters^50^— it has been hypothesized that such a disruption across the wider network may develop secondary to focal hippocampal damage and may not be directly caused by autoantibody-mediated mechanisms.^12^

In this context, a network view of LGI1-E also aligns well with the temporal dynamics of clinical manifestation, which typically involves a primary, often seizure-dominant symptom complex at onset and subsequently progresses to include additional features such as psychiatric symptoms and cognitive deficits with a latency of weeks to months.^42^ The long-term alterations of WM networks observed here thus suggest that LGI1-E may primarily cause focal brain damage initially and only subsequently affect other regions through their structural connections to this initial target – a process which could explain the latency and persistence of cognitive deficits.

### Implications for clinical trials

Given the prevalence of residual symptoms despite immunotherapy, it has been argued that LGI1-E and other forms of AE should be viewed as only partially immunotherapy-responsive conditions for which significant advances in clinical management are necessary to improve long-term outcomes.^1^ Consequently, objective markers that guide post-acute treatment are urgently needed because cognitive deficits are a key contributor to long-term disability^9, 18, 21^ and every fifth patient with LGI1-E lives in dependency.^51^ Our study presents the TDI as a novel neuroimaging marker that was highly sensitive to cognitive impairment in the post-acute stage and thus represents a promising new outcome measure for prospective clinical trials.

Furthermore, our approach explicitly captures patient-specific patterns of brain alterations because the TDI is agnostic to which particular regions show the strongest deviations from the reference group,^52^ yielding a highly personalized outcome marker. Not least, this comparison of individual MRIs to a healthy reference population represents the first step towards a normative neuroimaging framework – similar to diagnostic laboratory tests – in which the patient’s score is compared to a reference range and may thus serve as a biomarker to monitor disease course and treatment response.

### Limitations and Outlook

Limitations of our study include the cross-sectional design, which precludes inferences about the onset of network reorganization in LGI1-E and whether or not it is reversible. In this context, our study clearly shows that both structural brain changes and persistent cognitive deficits were not prevented by the applied immunotherapy. Therefore, our results call for future trials to evaluate if more aggressive or sustained immunotherapy is able to modify long-term disease course. Finally, while it is now clear that cognitive deficits represent a key outcome of LGI1-E, cognitive impairment shows a complex interrelationship with fatigue, which has recently been shown to impact patient-reported quality-of-life in the post-acute stage.^53^ Therefore, a natural next step will be to assess how brain network reorganization relates to persistent fatigue.

## Conclusion

Our study characterizes LGI1-E as a network disease that affects both limbic and extra-limbic brain systems and shows that a reorganization of structural brain networks is strongly linked to persistent and multi-domain cognitive impairment in the post-acute disease stage – despite immunotherapy and good overall recovery. These findings highlight the need for extended treatment strategies to improve long-term cognitive outcomes and propose a sensitive new neuroimaging marker to include in prospective clinical trials.

## Acknowledgments

This work was supported by the German Research Foundation (DFG), grant numbers 327654276 (SFB 1315), 504745852 (Clinical Research Unit KFO 5023 ‘BecauseY’), FI 2309/1-1 (Heisenberg Program) and FI 2309/2-1; and the German Ministry of Education and Research (BMBF), grant number 01GM1908D (CONNECT-GENERATE).

## Supplement

### Extended Methods

#### Participants

Between January 1^st^, 2013, and April 2^nd^, 2019, we prospectively enrolled 25 patients with anti-LGI1 encephalitis that presented to the Department of Neurology at Charité - Universitätsmedizin Berlin, Germany. Partial data of 16 patients have been analyzed in prior work,^11^ albeit regarding other imaging modalities. All patients fulfilled current diagnostic criteria^25^ and tested positive for anti-LGI1 antibodies in serum and/or CSF with indirect immunofluorescence assays. Using a standardized case report form, we systematically collected patient data including age, sex, symptoms at onset, MRI abnormalities, EEG abnormalities, malignant comorbidity, ICU treatment, follow-up time, relapse, treatment, and outcomes. As a control cohort, we included 25 healthy individuals who were matched to patients for age and sex, using the MatchIt package for R (version 3.0.2, https://github.com/kosukeimai/MatchIt). All participants gave informed written consent. The study was performed in accordance with the 1964 Declaration of Helsinki in its currently applicable version and was approved by the ethics committee of Charité - Universitätsmedizin Berlin.

#### Cognitive assessment

Cognitive performance in patients with anti-LGI1 encephalitis was assessed with standardized neuropsychological examination. Following previous approaches,^11^ test performance was quantified with composite scores across five cognitive domains known to be affected in AE: (1) visuospatial memory (Rey-Osterrieth Complex Figure test [ROCF];^54^ immediate and delayed recall scores), (2) verbal episodic memory (German edition of the Rey Auditory Verbal Learning Test [RAVLT];^55^ supraspan, interference, delayed recall scores), (3) attention (Test Battery for Attention Performance [TAP];^56^ median reaction times for tonic and phasic alertness and divided visual and auditory attention), (4) executive functions (TAP Go/NoGo median reaction time), and (5) working memory (forward and backward digit span test scores^57^). Where applicable, composites were calculated by first *z*-scoring the individual sub-scores and subsequently averaging over these *z*-scores within a cognitive domain. In this process, *z*-scores of reaction times were inverted to ensure that higher composite scores universally indicate better performance across all cognitive domains.

#### MRI acquisition

Brain MRI was performed at the Berlin Center for Advanced Neuroimaging (BCAN), using a 3 Tesla Tim Trio scanner (Siemens, Erlangen, Germany) equipped with a 12-channel phased array head coil. For DWI acquisition, a single-shot echo planar imaging sequence was used (repetition time, [TR] = 7500 ms; echo time [TE] = 86 ms; field of view [FOV] = 240 x 240 mm^2^; voxel size = 2.5×2.5×2.3 mm^3^, 61 slices, 64 non-colinear directions, b-value = 1000 s/mm^2^, one b=0 image). For anatomical scans, a volumetric high-resolution T1-weighted magnetization prepared rapid acquisition gradient echo (MPRAGE) sequence was performed (TR/TE/inversion time [TI] = 1900/2.55/900 ms, FOV = 240 x 240 mm^2^, matrix size = 240 x 240, 176 slices, slice thickness = 1 mm).

#### MRI preprocessing and connectome construction

DWI preprocessing and tractography were performed using MRtrix3,^28^ FSL,^29^ and Advanced Normalization Tools^30^ as described previously.^32^ In summary, we performed denoising, eddy-current correction, and motion correction for DWI images. To achieve global intensity normalization, we conducted bias field correction and DWI group normalization.

Anatomical T1-weighted scans were parcellated into 84 cortical and subcortical regions of interest (ROI) using the Desikan Killiany atlas^31, 58^ and segmented into white matter (WM) and gray matter (GM) using FSL FAST.^59^ Then, T1-weighted scans were registered to DWI images and, using a group average response function for normalization, fiber orientation density functions (ODFs) were obtained using single-shell, single-tissue constrained spherical deconvolution. Anatomically constrained probabilistic tractography (ACT) and spherical-deconvolution informed filtering of tractograms 2 (SIFT2) was conducted to create biologically accurate connectomes from 2 x 10^7^ streamlines.^26, 32–34^ The output file was an 84 x 84 structural connectivity matrix, where each cell value represents the absolute streamline count of the respective ROI-to-ROI connection. To achieve inter-participant connection density normalization, each value of the matrix was multiplied by the participant-specific fiber density SIFT2 proportionality coefficient (μ), resulting in weighted and normalized structural connectivity matrices.

### Region-to-network mapping

Assignment of anatomical regions to functional brain systems was implemented with a custom mapping algorithm. To this end, each of the 84 cortical and subcortical regions in the Desikan Killiany atlas was assigned to one of the seven canonical resting-state networks from the Multiresolution Intrinsic Segmentation Template (MIST7).^38^ The MIST7 atlases include subcortical and cortical brain regions which are grouped into seven functional clusters including the cerebellum, mesolimbic network, somatomotor network, visual network, default mode network, fronto-parietal / visual downstream network, and ventral attention network / salience network / basal ganglia / thalamus. The MIST7 atlases were resampled to 1 mm^3^ isotropic resolution using the Statistical Parametric Mapping 12 software ( http://www.fil.ion.ucl.ac.uk/spm/software/spm12/) to match the resolution of the Desikan Killiany atlas. Both atlases were provided in Montreal Neurological Institute (MNI) standard space. For each ROI in the structural atlas, a binary nifti file was created, and the number of voxels overlapping with each of the binary MIST7 network niftis was calculated. The functional assignment for each structural ROI was then given by the functional MIST7 system which showed the maximum number of overlapping voxels with the anatomical region. Finally, these programmatically assigned functional labels were quality-controlled by visual inspection.

#### Network analysis and graph-theoretical measures

The topological characteristics of individual WM networks —commonly referred to as structural ‘connectomes’^60^— were estimated with custom code and subroutines from the Brain Connectivity Toolbox (BCT, www.brain-connectivity-toolbox.net).^35^ To this end, symmetrical connectome matrices were constructed from the tractography results, where the matrix diagonal was set to zero to discard self-connections. Furthermore, individual connectomes were normalized by the maximally observed connectivity strength, such that connection strengths were standardized to lie in [0, 1] for each participant. Subsequently, network sparsity was implemented by thresholding the connection strength and setting all elements below the respective threshold to 0, following common practice.^23, 35^ Therein, we applied a set of linearly increasing cut-off values from 0 to the 95th percentile by increments of 5, resulting in 20 thresholded connectomes of increasing sparsity for each participant. Since it is an open question which sparsity threshold is optimal for subsequent network analyses (and since this threshold is arbitrary by definition), we applied a summary procedure that is agnostic to the particular sparsification thresholds. To this end, the respective network measures were calculated for each individual threshold and then aggregated as the area-under-the-curve (AUC) over all 20 thresholds, yielding a single summary measure.

To achieve this aggregation step, individual thresholded and normalized connectomes were subjected to an analysis of network topology for weighted and undirected graphs, in which we estimated the node degree (ND) and betweenness centrality (BC) for every brain region. The mathematical formulation and conceptualization of these graph-theoretical measures for brain networks have been described in detail in previous work^23, 35, 61^. Briefly, ND quantifies the number of structural links that pass through a given node, yielding a measure of its overall ‘connectedness’. Note that ND values here were symmetrical since network graphs were undirected, such that the connectivity between region A ➔ region B was identical to region B ➔ region A. Moreover, BC values for weighted networks were calculated for every brain region. This measure expresses the proportion of all shortest paths in the network that pass through a given node, commonly interpreted to indicate its ‘hubness’ (that is, its relative importance in the network) as it expresses the propensity of the node to exert a ‘bridging role’ in the network.^23, 35, 39^

#### Topology deviation index (TDI)

To quantify changes in the whole-brain topology of structural connectomes, we defined a topology deviation index (TDI) which rests on recent methodological developments^52^ and was computed as follows: First, a reference topology was calculated over all healthy participants as the median BC value for each brain region, yielding a 1×84 vector of reference values for the Desikan-Kiliany atlas. Herein, we chose BC as the target graph metric because it expresses topological network organization rather connectivity alone and because prior group analyses suggested wide-spread and bidirectional connectome reorganization in patients with LGI1-E (**Fig. 2**). In a next step, we then related this reference distribution to the regional centrality values of individual participants. Topological deviation was subsequently calculated as *TDI* = 1 − *ρ*_*S*_, i.e., the non-parametric correlation distance between the reference centrality values and the individual centrality values, where *ρ*_*S*_ refers to Spearman’s rank correlation coefficient. Here we chose rank-based over product-moment correlation because the former does not assume a linear relationship between the two variables and is less sensitive to outliers. The TDI thus resolves to zero if an individual brain topology perfectly adheres to the reference topology in rank space and gradually increases to >0, the more the individual values deviate from this reference (see **Fig. 4B**). Note that this approach thus captures topological network changes across all brain regions simultaneously, yielding a single whole-brain deviation value for each participant.

#### Statistical analyses

Univariate between-group differences were assessed as the standardized effect size for independent-sample t-tests (Cohen’s *d*), unless otherwise stated. The ordinal mRS and CASE outcomes scores in Figure 3 were assessed once at peak illness and once at post-acute follow-up for each patient, such that these samples were considered paired and assessed with the nonparametric signed rank test. Continuous relationships were tested with product-moment correlation or rank-based correlation, as indicated. Differences in network topology between patients and controls were assessed with a permutation approach. The null hypothesis under this regime posits that there is no between-group difference and that, consequently, there is no effect of whether a particular participant is labeled a patient or a control. To test this hypothesis, one then estimates the null distribution of the statistic under question by randomly permuting the group labels and recalculating the statistic for every permutation. The ensuing *p*-value of this test is then given as the number of instances in which the randomly permuted statistics surpass the value of the empirically observed statistic, divided by the total number of random permutations (here n = 5000). When a variable was repeatedly tested against multiple other measures, we applied the Benjamini-Hochberg procedure to control the false discovery rate (FDR).^37^

#### Data Availability

Anonymized data can be shared by the corresponding authors upon reasonable request from qualified investigators. Analysis code underlying the results of the current study will be made publicly available upon acceptance of the manuscript.

## Notes

### Competing Interest Statement

The authors have declared no competing interest.

